# scPipe: An extended preprocessing pipeline for comprehensive single-cell ATAC-Seq data integration in R/Bioconductor

**DOI:** 10.1101/2023.09.25.559230

**Authors:** Shanika L. Amarasinghe, Phil Yang, Oliver Voogd, Haoyu Yang, Mei R. M. Du, Shian Su, Daniel V. Brown, Jafar S. Jabbari, Rory Bowden, Matthew E. Ritchie

## Abstract

`scPipe` is a flexible R/Bioconductor package originally developed to analyse platform-independent single-cell RNA-Seq data. To expand its preprocessing capability to accommodate new single-cell technologies, we further developed `scPipe` to handle single-cell ATAC-Seq and multi-modal (RNA-Seq and ATAC-Seq) data. After executing multiple data cleaning steps to remove duplicated reads, low abundance features and cells of poor quality, a *SingleCellExperiment* object is created that contains a sparse count matrix with features of interest in the rows and cells in the columns. Quality control information (e.g. counts per cell, features per cell, total number of fragments, fraction of fragments per peak) and any relevant feature annotations are stored as metadata. We demonstrate that `scPipe` can efficiently identify “true” cells and provides flexibility for the user to fine-tune the quality control thresholds using various feature and cell-based metrics collected during data preprocessing. Researchers can then take advantage of various downstream single-cell tools available in Bioconductor for further analysis of scATAC-Seq data such as dimensionality reduction, clustering, motif enrichment, differential accessibility and cis-regulatory network analysis. The `scPipe` package enables a complete beginning-to-end pipeline for single-cell ATAC-Seq and RNA-Seq data analysis in R.

## Introduction

Single-cell sequencing technology has undergone rapid development in the past decade to allow researchers to study cellular heterogeneity across multiple omic modalities, including the transcriptome, epigenome and proteome. Single Cell Assay for Transposase Accessible Chromatin Sequencing (scATAC-Seq), is a relatively recent approach for profiling chromatin accessibility at single-cell resolution (1) that has been widely used to define chromatin state across cell types, discover cis- and trans-regulatory regions, identify master regulators, and characterise gene regulatory networks (2). The four main protocols for scATAC-Seq include the combinatorial indexing approach (sci-ATAC-Seq) (3), microfluidics-based methods (scATAC-Seq) (1), nano-well based protocols (*µ*scATAC-Seq) (4) and droplet-based (10X scATAC-Seq, dscATAC-Seq and dsciATAC-seq) approaches (5, 6). The general workflow for scATAC-Seq data analysis comprises of [1] preprocessing: demultiplexing, adaptor trimming, read mapping, quality control, cell calling and multiplet removal *(optional)*; [2] feature matrix construction: defining regions via peak calling or genome binning, counting defined features, transformation and dimensionality reduction; and [3] downstream analysis: cell clustering, secondary peak calling, visualisation, differential accessibility analysis and cis-regulatory network analysis (7, 8).

The growing popularity of scATAC-Seq technology warrants the development of analysis tools that can preprocess this type of data efficiently and effectively. There are currently 14 tools that can preprocess scATAC-Seq data that are summarised in Table 1. Some are written in R (i.e. `ArchR`, `BROCKMAN`, `ChromSCape`, `chromVAR`, `Destin`, `scABC`, `SCRAT`, `Snaptools`) while others are Python and Shell based. Furthermore, the feature matrix construction step varies amongst these tools where some of them use bulk peak calls, TSS regions, *k*-mers, or a genome binning approach. It has been shown previously that out of these, a genome binning approach is more sensitive to the detection of rare open chromatin regions present in a sub-population of cells (9).

**Table 1.** Summary of current packages available for scATAC-Seq data preprocessing.

| Method / Tool Name | Year | Language | Input | Multiple Technologies? | Feature Matrix Construction Method |
| --- | --- | --- | --- | --- | --- |
| APEC | 2019 | Python | *FASTQ*, matrix + peaks | ✓ | Peak |
| ArchR | 2020 | R | *BAM*, fragments | ✓ | Bin |
| BROCKMAN | 2018 | R | *FASTQ* | × | *k*-mer |
| Cell Ranger ATAC | 2018 | Command line | *FASTQ* | × | Peak |
| ChromScape | 2019 | R Shiny | Various | × | Bin, Peak, TSS |
| chromVAR | 2017 | R | *BAM* + peaks | × | Motif, *k*-mer |
| Cusanovich *et al.* 2018 | 2015 | Scripts | *BAM* | × | Peak |
| Destin | 2019 | R | *FASTQ*, *BAM* + peaks | ✓ | Peak |
| Gene scoring | 2019 | Scripts | *FASTQ* | × | TSS |
| scABC | 2018 | R | *FASTQ* | × | Peak |
| scasat | 2018 | JupyterNotebook | *BAMs* | × | Peak |
| scATAC-pro | 2019 | Command line | *FASTQ*, various | ✓ | Peak |
| SCRAT | 2017 | R,WEB | *BAMs* | × | Various |
| Snaptool | 2019 | Python | *FASTQ* | ✓ | Bin, Peak |

A current limitation of these tools is that most cannot handle data from multiple technologies (i.e. droplet-based and plate-based). Furthermore, most of avaialble tools require the use of specific data structures that have limited compatibility with current well-known, light-weight, single-cell data structures such as *SingleCellExperiment* objects (*SCEs*) in R/Bioconductor.

In this study, we extend `scPipe` (10) to enable handling of scATAC-Seq data and generation of *SCE* objects that can be manipulated using various *SCE*-friendly downstream analysis tools available in R/Bioconductor (11). An *SCE* container, most popular for scRNA-Seq data storage, can easily be used to store scATAC-Seq data due to its flexibility (i.e., rows representing features (genomic regions) and columns representing cells). `scPipe` is able to take scATAC-Seq reads (*FASTQ* format) as input, which are demultiplexed (if needed) and filtered based on quality and Ns (i.e. non-called bases), aligned to the reference genome and filtered based on various quality metrics such as mapping rate, fraction of reads mapping to the mitochondrial genes and the number of duplicate or high-quality reads. Cleaned up data will then be used to conduct cell calling and optionally peak calling, to generate the final feature by cell sparse matrix which is stored as a *SCE* object. These improvements also allow `scPipe` to be used for multiome projects that collect both scRNA-Seq and scATAC-Seq on the same cells.

## Materials and Methods

### Architecture of the scPipe scATAC-Seq module

`scPipe` was developed using the R (12) / Bioconductor (13) platform, with the underlying code written in C++, R and Python with the `Rcpp` (14) and `reticulate` (15) packages used to wrap the C++ and Python code for R, respectively. Data from both UMI (Unique Molecular Identifiers) and non-UMI protocols can be handled by `scPipe`. The pipeline for scATAC-Seq data preprocessing is initiated with *FASTQ* files and outputs include a feature count matrix and a variety of quality control (QC) statistics and a standalone HTML report generated using `rmarkdown` (16) that contains a summary of the QC statistics collected during data preprocessing.

### Demultiplexing and read alignment

Demultiplexing is performed by the *sc_atac_trim_barcode()* function (Figure 1A) using a similar approach to that used in `scPipe` for scRNA-Seq analysis. The scATAC-Seq module can accommodate data in .*f astq* format as well as .*csv* format, with the user defining which format the data is in and the relevant demultiplexing strategy will be executed. In brief, the barcode is either extracted from the reads themselves based on the entries in a *csv* file where the second column of the file contains the barcodes or from the sequences of a complementary *FASTQ* file and appended to the read names in the *FASTQ* files containing the ATAC sequences. A read correction step is incorporated to ensure sequencing errors that appear in barcodes are identified and removed to avoid unwanted data loss. In summary, after the barcodes are determined, error correction is carried out with a hamming distance of 1 to correct for sequencing errors. Each barcode can be optionally vali-dated against a known list of valid barcodes (either as is or the reverse complement) and the frequency of each valid barcode is then counted. `scPipe` corrects barcodes that are not matched to the valid barcode list by allowing for one mis-matched base in comparison with the valid barcode list, and counting the number of corrected barcodes in the complete dataset. In the case that multiple sequences are found matching the valid barcode list with one mismatched base, each mismatched base’s quality (Q) score is compared and the sequence with the lowest Q score at the mismatch position is considered valid. Demultiplexed data is stored in 4 data files; *i*.*e*. complete-match: no error correction was necessary for the data in this file; partial-match: error-corrected reads; no-match: data that were unable to be demultiplexed even after error correction; and full-data: a concatenated version of the data across the 3 aforementioned categories. The user is able to select either one of the more stringent files or the complete dataset for downstream analysis (by default, the complete dataset is used).

**Fig. 1.**
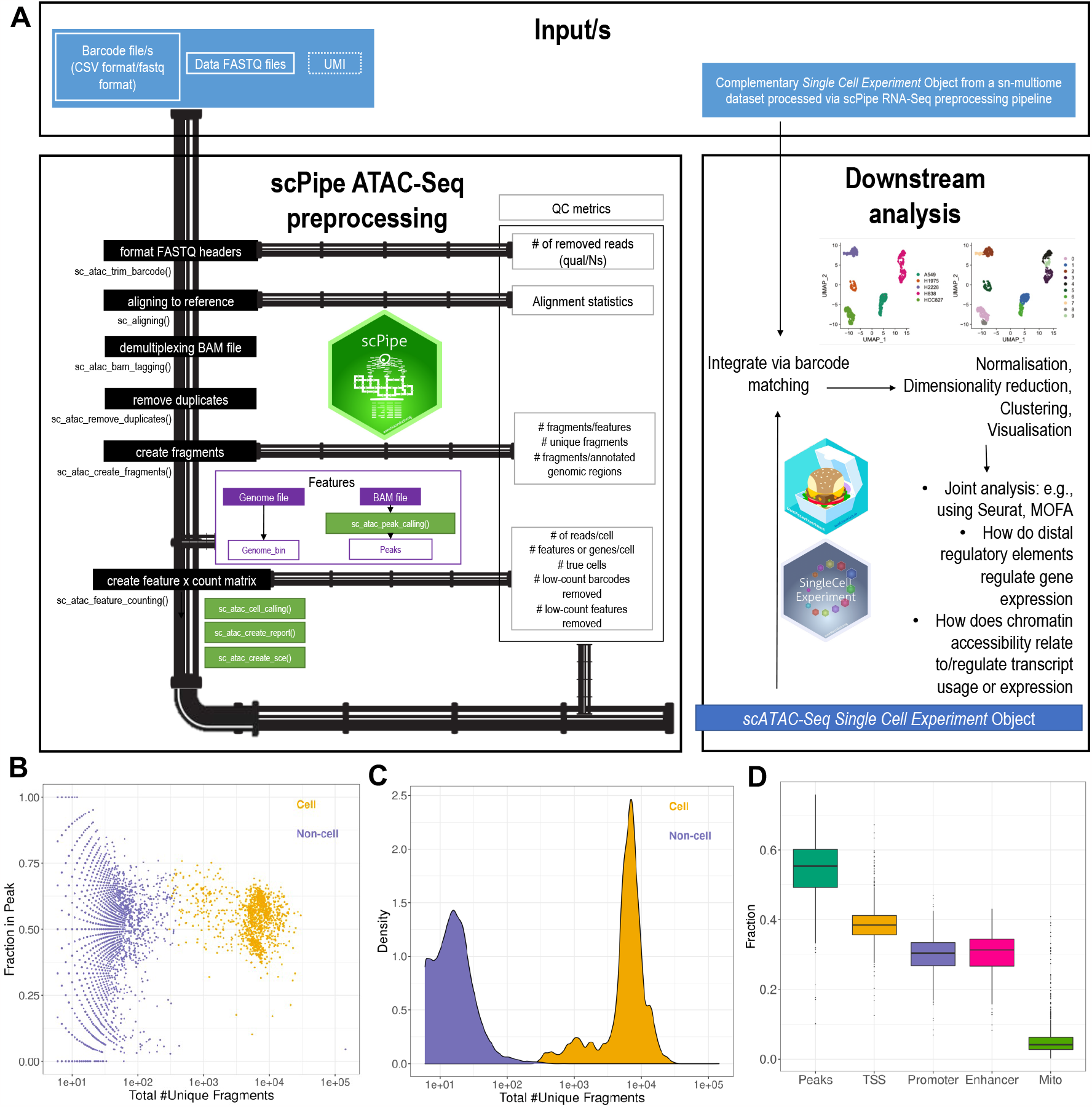
Overview of the scPipe scATAC-Seq module and its QC outputs. (A) The pipeline is shown on the left and the QC metrics gathered during preprocessing are shown on the right. Purple coloured boxes denote the inputs and light colour purple define the inputs that are optionally accepted as they are not incorporated into current scATAC-Seq library preparations yet. Blue colour box depicts the final output. Black boxes denote the main pipeline steps that should be followed. Green boxes denote the steps that are running within these the main pipeline without having to call them specifically, still can also be called separately if needed. (B) QC plot showing the separation of “cell” and “non-cell” based on the fraction of fragments overlappping peaks (y-axis) vs total number of fragments (x-axis) after cell calling step of scPipe. (C) QC plot showing the separation of cell and “non-cell” based on read density (y-axis) vs total number of fragments (x-axis) after cell calling step of scPipe. (D) QC plot showing the fraction of features overlapping different functional regions (i.e., Peaks, TSS, Promoter, Enhancer, Mitochondrial genes).

Read alignment is run with the function *sc_aligning()* which uses *Rsubread::align()* (17) internally (Figure 1A). The user has to define the technology (i.e. RNA or ATAC) to execute the most appropriate alignment approach (i.e. single-end alignment for RNA and paired-end for ATAC). The resulting *BAM* file contains the read name in the first column, and unmapped reads are denoted by an asterisk (*∗*).

### Demultiplexing aligned reads, removing duplicates, creating fragments and peak calling

Demultiplexing aligned reads is performed by the *sc_atac_bam_tagging()* function which extracts the barcode information from the read names of the aligned *BAM* file and generates a new column with the tag CB:z: to record the cell barcode (Figure 1A). As some reads may not have an assigned barcode, this column in the resulting demultiplexed *BAM* file may be empty in such positions. The main difference in the scATAC-Seq module compared to original scRNA-Seq module is the need to extract biological information from both reads in the former. Therefore, it was important to retain both reads (i.e.forward and reverse) for scATAC-Seq data as compared to retaining only the forward of the pair of reads for scRNA-Seq module where reverse will be the barcode followed by the polyA tail, which can be discarded after demultiplexing of the read takes place.

Removal of duplicate reads uses `samtools`, where the *BAM* file is processed with the *removeduplicates* function (18). This step can be run externally to `scPipe` and the resulting *BAM* file can be re-entered to the `scPipe` pipeline if `samtools` is not installed locally.

Fragment file generation was adopted from a python-based tool named `Sinto` (version 0.8) (19). Briefly, the fragment file is in .*BED* format created from the aligned read file (i.e. *BAM*) with the position of each Tn5 integration site, barcode of the cell that the fragment belongs to and the number of times the fragment was sequenced while collapsing the PCR duplicates. This is achieved by first extracting the cell barcode sequence associated with each read and adjusting the alignment positions for the 9 bp Tn5 shift by applying +4/-5 to the start and end position of the paired reads. Next, fragments below a certain quality threshold and a size larger than a maximum provided by the user are removed before collapsing any duplicates (if present). This python-based capability was integrated into the `scPipe` R package using `basilisk` (20).

Feature matrix construction can be via a .*bed* file provided to the workflow (e.g. bulk peak file from `MACS3`) or using the reference genome (i.e. ‘`genome_bin`’ approach). In the ‘`genome_bin`’ approach, the genome is cut in to chunks of user-defined size and the overlap is calculated between the *bed* file and the fragment file. Using such an approach is resource intensive, yet more sensitive to rare open regions than using a bulk approach. Furthermore, being able to converge on the same features via this approach will make downstream integration of the multiple scATAC-Seq data sets more convenient. An optional peak calling step can also be carried out using the R version of `MACS3` (`MACSr` (21)) which has been implemented to execute bulk-level peak calling across all the data. This is a faster method than the in-built *genome −bin* approach described above to identify features to generate the feature *×* cell matrix.

### Generating the feature *×* cell matrix, SCE objects and QC statistics

The externally created (e.g. via `Cell Ranger`) or `scPipe` generated feature matrix and fragment file is used as the main input to generate the feature *×*cell count matrix. This step involves multiple filtering steps including cell calling and row-wise (i.e. feature-level), and column-wise (i.e. cell-level) filtering as well as read correction for the Tn5 cut site. A *GenomicAlignments* object (22) is generated for the fragments with cell barcode information and the feature file (peaks/bins) with the feature coordinate information which is then overlapped to generate the unfiltered counts matrix. This matrix then enters the cell calling step to distinguish “true” cells from ambient background DNA. It has been previously been shown that filtering cells based on multiple QC metrics is the most effective approach to do cell calling (7), and this approach has been adopted in `scPipe`. This can be called with the parameter *cell_calling=filter* within the *sc_atac_feature_counting()* function. The resulting matrix is then converted to a binary matrix as well as a *SCE* object (11). An HTML report is generated using `rmarkdown` as an optional output of this step. This report summarises the QC metrics gathered during preprocessing to allow data quality to be visually assessed (Figure 1B-D), and is easy to share with collaborators. `ScPipe`’s ATAC-Seq module also logs all details in the workflow so that a user can track the progress if a step fails or wants to further explore the pipeline parameters used.

### Integration of SCE objects

Another important aspect of `scPipe` is the ability to integrate datasets where a common barcode file is available to match the cells between *SCE* objects to create a combined *SCE* object, as occurs for example in 10X Genomics multiome experiments. The R/Bioconductor package `MultiAssayExperiment` is used for this (23) to output combined *SCE* objects with `colData()` from the same barcodes bound together (Figure 1A). If there are barcodes with missing data from one or more *SCE* objects, they will still be included with the addition of ‘NAs’ to columns corresponding to the cells that do not share common barcodes. Storing data in this way allows the user to further leverage Bioconductor software and design principles to make biologically relevant inferences (e.g. to correct for batch effects, perform integration and clustering).

### Human lung adenocarcinoma cell line dataset

The cell culture and sample preparation of mixtures of cells from different cell lines was performed as previously described (24). Briefly, five human lung adenocarcinoma cell lines (A549, H1975, H2228, H838 and HCC827) were obtained from ATCC (https://www.atcc.org/) and cultured separately in Roswell Park Memorial Institute (RPMI) 1640 medium with 10% fetal calf serum and 1% penicillin-streptomycin at 37°C with 5% carbon dioxide until near 100% confluency. Cells were counted manually using a hemocytometer and mixed with equal number to form single-cell suspension with around 2,000,000 cells. Cells were permeabilised with nuclei EZ Lysis Buffer from Sigma-Aldrich supplemented with Protector RNase Inhibitor and filtered to isolate nuclei. Nuclei underwent fluorescence-activated nuclei sorting (FANS) and 1,600,000 nuclei were output from the sorter. After concentrating the nuclei suspension to achieve 10,000 nuclei recovery during Gel Bead-In EMulsions (GEM) generation, 11,960 nuclei were input to form GEMs that were used to generate 10X Multiome (GEX + ATAC) libraries. The cDNA library was sequenced using Illumina NextSeq 500 with recommended cycles and the ATAC library was sequenced using Illumina NextSeq 500 with the custom recipe. FASTQ files were then generated with `Cell Ranger ARC` 2.0.0 *mkfastq*. The raw data are available from GEO under accession number GSE224045.

### scATAC-Seq data preprocessing

We have processed the scATAC-Seq data in two ways; 1) via the `Cell Ranger ATAC` pipeline, and 2) via the `scPipe` pipeline. The scRNA-Seq data was processed through the standard `Cell Ranger` pipeline. The paired read were aligned to the hg38 reference. The tool `demuxlet` (25) was used to assign the cell line identity (i.e., ground truth) to the data using variant information as previously described (24). Next we developed custom scripts (available via GitHub) using mainly `ggplot2`(26), `Seurat` (27) and `NMI` R packages to generate plots for visualisation and comparison.

## RESULTS

### Resource requirements of scPipe on scATAC-Seq data

For our data containing 112,463,635 reads, the pipeline required approximately 604 minutes (10 hours) to complete and 540 GB of RAM allocated. The longest processing step was *sc_aligning*, which internally uses `Rsubread`, and required 78 minutes to complete.

### Downstream comparison between scPipe and Cell Ranger ATAC

We compared the clusters identified from `scPipe` preprocessed data to those obtained from the `Cell Ranger ATAC` pipeline, a popular tool for 10X scATAC-Seq preprocessing. A dataset that contains around 5,800 cellswith ground truth available in the form of cell line identity labels obtained using variant information was used to compare the results from the two approaches (Figure 2A). `Cell Ranger` returned 5,728 cells and `scPipe` 5,599 cells after QC, with 5,267 in common (Figure 2B). Inspection of features overlapping the mitochondrial genes as a QC metric shows that `Cell Ranger` has called more dead/poor quality cells that should be excluded from downstream analysis (Figure 2C). On the other hand, the mitochondrial contamination is lower and similar for `scPipe`-only called cells and those identified by `scPipe` in common with `Cell Ranger` (Figure 2C’). Overall, there was high concordance between the called cell (Figure 2D) and feature counts (Figure 2D’) within for the cells in common between the two pipelines. Moreover, “`scPipe`-only” called cells contained counts and features that were more similar those found in common than the counts and features from “`Cell Ranger`-only” called cells, whose distributions were concentrated towards the lower ranges for both quantities. The UMAP (28) generated from the `Cell Ranger` output shows that the 129 cells that appear in the `Cell Ranger` results but not in `scPipe` tend to cluster together away from the large clusters or in the margins of them (Figure 2E). In contrast, the UMAP generated from the `scPipe` output shows that the cells that only appear in `scPipe` results but not in `Cell Ranger` tend to be located within the main clusters (Figure 2E’). Comparison of the cell line labels assigned to these cells using `demuxlet` (25) to the clustering results obtained using the scATAC-Seq data alone in a `Seurat` analysis, was made by calculating the adjusted rand index (ARI) and normalised mutual information (NMI) scores which should be closer to 1 when the clustering is highly concordant between the two approaches. For the `Cell Ranger` data, an ARI of 0.56 and NMI of 0.64 (Figure 2F) were obtained. The `scPipe` generated output resulted in a higher ARI of 0.68 and NMI of 0.75 (Figure 2F’), indicating better concordance between the `Seurat` clustering and `demuxlet` results. Taken together, `scPipe` seemed to identify higher quality cells that were better separated according to the biological signal present in the data relative to the `Cell Ranger` processed data.

**Fig. 2.**
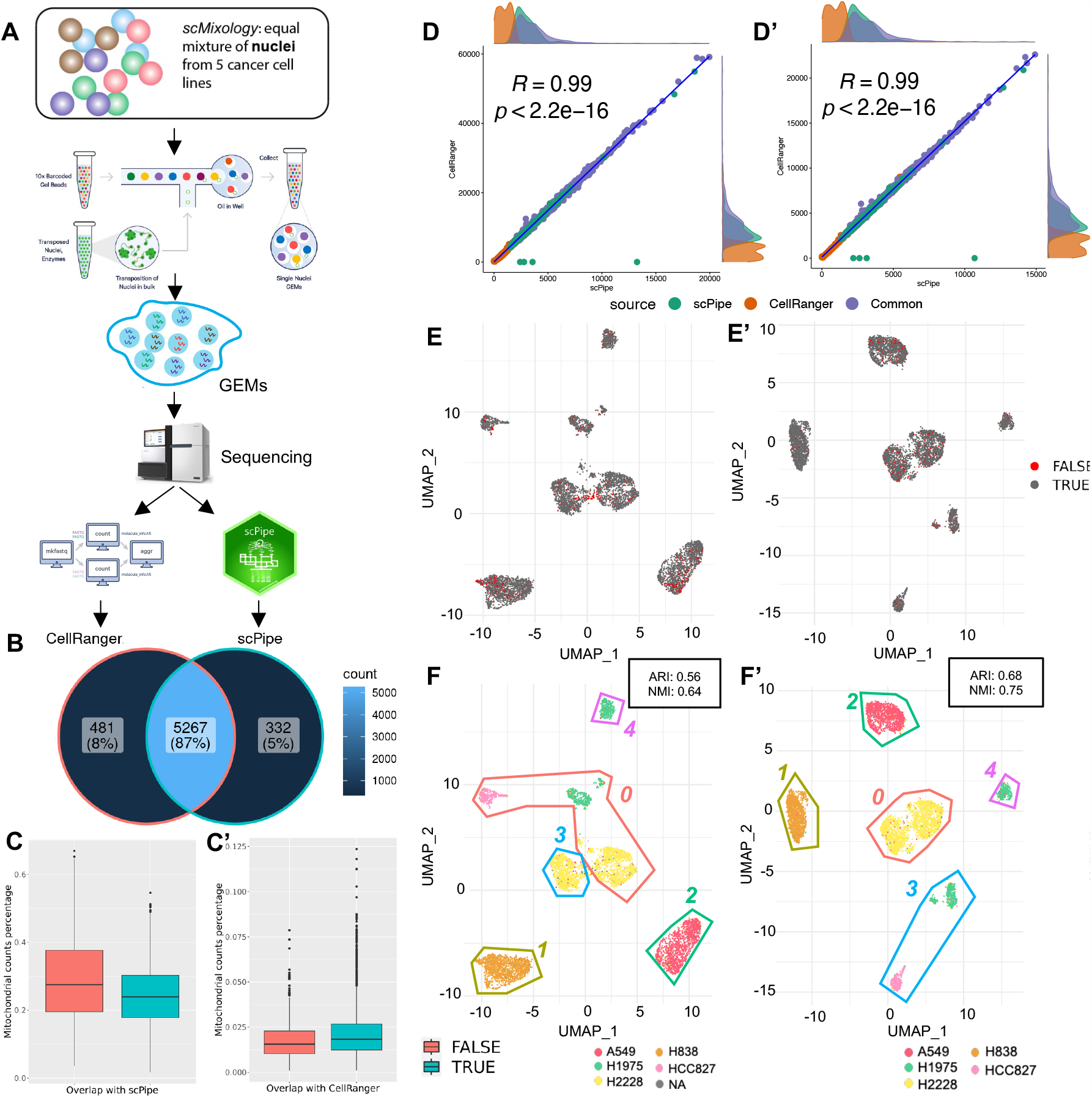
Comparing scPipe and Cell Ranger on a 10x dataset. (A) Summary of experimental design, which used cells from 5 distinct lung adenocarcinoma cell lines. An equal mixture of cells and nuclei were captured by the 10X protocol, and sequenced on the Illumina platform (See Materials and Methods). *FASTQ* files were generated by `Cell Ranger ARC` 2.0.0 and these reads were processed by `scPipe` and `Cell Ranger`. The panels of this figure pertain to the 80% results. (B) Venn diagram showing the overlap of cell barcodes detected by `Cell Ranger` and `scPipe`. (C) Box plot showing the percentage of mitochondrial gene counts in cells that are called by `Cell Ranger` and are common with/unique in comparison to `scPipe` or (C’) `scPipe` and are common with/unique in comparison to `Cell Ranger` output. (D) Scatter plot of the per cell total counts and (D’) number of features per cell obtained from `Cell Ranger` and `scPipe` in cells called in common between the two and cells that were unique to each of the two pipelines. Marginal density plots show the count distributions for each category. (E) UMAP plot generated for `Cell Ranger` and (E’) `scPipe` output. Cell barcodes that only exist in (E) `Cell Ranger` or (E’) `scPipe` are highlighted in red. (F) UMAP coloured by the ground truth for `Cell Ranger` and (F’) `scPipe`. `Seurat` identified clusters are demarcated by coloured lines and numbered. The ARI and NMI values calculated per dataset are shown in the top right of the panels.

## Discussion

To ensure flexibility, `scPipe` can preprocess data from a variety of different scATAC-Seq protocols that includes droplet-based and plate-based methods. `scPipe` can also handle combinatorial barcoding options when demultiplexing the reads, and can handle Unique Molecular Identifier (UMI) based scATAC-Seq approaches if they arise in the future. The resulting preprocessed data output by `scPipe` is stored in a versatile *SingleCellExperiment* (*SCE*) format, which means that the data can be further analysed downstream using *SCE*-object friendly tools available in R/Bioconductor. We demonstrate that data preprocessing with `scPipe`’s scATAC-Seq preprocessing workflow produces comparable results to those from 10X’s `Cell Ranger ATAC` pipeline. As the integration of numerous large-scale data modalities becomes more routine, `scPipe` provides convenient and scalable preprocessing options for both scRNA-Seq and scATAC-Seq data, allowing data integration and joint analyses.

## Availability

`scPipe`’s scATAC-Seq preprocessing module is available from Bioconductor (https://bioconductor.org/packages/scPipe) and code used for the timing calculations and generating Figure 2 (panels B-F) are available from GitHub (https://github.com/shaniAmare/scPipe-Manuscript-Materials).

## ACKNOWLEDGEMENTS

This work was supported by funding from the Chan Zuckerberg Initiative DAF, an advised fund of Silicon Valley Community Foundation (Grant No. 2019-002443 to M.E.R.), Australian National Health and Medical Research Council (NHMRC) Investigator Grant (2017257 to M.E.R.), the Australian Research Council (Discovery Project No. 200102903 to M.E.R.), the Genomics Innovation Hub, Victorian State Government Operational Infrastructure Support, Australian Government NHMRC IRIISS and support from the Australian Cancer Research Foundation. The authors are grateful to Dr Saskia Freytag and Mr Reza Ghamsari for providing feedback on this manuscript.

